# Ozone sensitivity of diverse maize genotypes is associated with differences in gene regulation, not gene content

**DOI:** 10.1101/2021.05.06.442991

**Authors:** Adalena V. Nanni, Alison M. Morse, Jeremy R. B. Newman, Nicole E. Choquette, Jessica M. Wedow, Zihao Liu, Andrew D. B. Leakey, Ana Conesa, Elizabeth A. Ainsworth, Lauren M McIntyre

## Abstract

The maize pangenome has demonstrate large amounts of presence/absence variation and it has been hypothesized that presence/absence variation contributes to stress response. To uncover whether the observed genetic variation in physiological response to elevated ozone (a secondary air pollutant that causes significant crop yield losses) concentration is due to variation in genic content, and/or variation in gene expression, we examine the impact of sustained elevated ozone concentration on the leaf tissue from 5 diverse maize inbred genotypes (B73, Mo17, Hp301, C123, NC338). Analysis of long reads from the transcriptomes of the 10 conditions found expressed genes in the leaf are part of the shared genome, with 94.5% of expressed genes from syntenic loci. Quantitative analysis of short reads from 120 plants (twelve from each condition) found limited transcriptional response to sustained ozone stress in the ozone resistant B73 genotype (151 genes), while more than 3,300 genes were significantly differentially expressed in the more sensitive NC338 genotype. The genes underpinning the divergence of B73 from the other 4 genotypes implicates ethylene signaling consistent with some findings in Arabidopsis. For the 82 of the 83 genes differentially expressed among all 5 genotypes and the 788 of 789 genes differentially expressed in 4 genotypes (excluding B73) in sensitivity to ozone is associated with oxidative stress tolerance being associated with a weaker response to a reactive oxygen species (ROS) signal and suggests that genetic variation in downstream processes is key to ozone tolerance.

## Background

Tropospheric ozone is a prevalent, phytotoxic air pollutant, which significantly reduces global crop production [1–3]. In the U.S. ozone pollution over the past 30 years was estimated to decrease maize yields by 10% [2]; a loss equivalent to the impact of aridity stress, temperature stress or nutrient stress [3]. Exposure of plants to elevated ozone concentration causes reactive oxygen species (ROS) production in cells and knowledge of the responses this elicits is expected to aid understanding of oxidative stress associated with other abiotic and biotic factors [4, 5]. Loss of photosynthetic capacity associated with accelerated senescence is a key response of maize to elevated ozone and differs among genotypes [6]. Ozone stress increased the heritability of photosynthetic traits and strengthened genetic correlations among traits in maize with substantial differences among hybrids observed [7].

Within maize accessions there has been discussion of the divergence of genomes [8] with reports of a megabase deletion [9] as well as reports of pervasive copy number, presence/absence variation [10] and exon shuffling [11]. However, large portions of grass genomes are syntenic [12–16]. A recent *de novo* assembly of Mo17 [17] and corresponding in depth comparison to B73 found only 122 genes present exclusively in one of the two genomes, concluding that the sub-genomes were nearly identical. Despite this macrocollinearity, conservation of genes and gene order, there are differences in genic rearrangements, such as insertions, deletions, amplifications, inversions, and translocations [reviewed in 18].

The contribution of the syntenic genome to gene expression in leaf tissues among diverse maize genotypes, and the degree of conservation of response to ozone stress is unknown and is an important component in understanding oxidative stress response. This study represents an opportunity to investigate the genetic basis of physiological response of maize to ozone stress. Differences in ozone response among maize genotypes could be due to structural variation in the genomes themselves or to differential regulation of expression in a common set of genes. To test these possibilities, we sequenced the leaf transcriptome of five diverse maize genotypes, B73, Mo17, C123, NC338 and Hp301, in ambient and elevated ozone conditions. While C123 and Mo17 are somewhat closely related, the other genotypes are divergent, and Hp301 is a popcorn [See Supplementary Figure 1 in 19]. These five genotypes are selected based on variation in physiological response to ozone in field experiments [7] where B73 was the least sensitive of the 5 genotypes. We use long read sequencing to accurately identify and compare the set of transcripts expressed in each genotype and condition, as well as to understand the potential contribution of differential splicing to the stress response. We collected leaf tissue from each of the 5 genotypes in each condition (n=10) and assemble transcriptomes for each condition separately. These transcriptomes are combined and we use the combined transcriptome to estimate gene expression and test for differences in response to ozone, for each of the 5 genotypes using samples from a controlled greenhouse experiment with twelve independent replicate plants for each genotype/condition (n=120).

Our results indicate that the qualitative composition of the leaf transcriptome is conserved across maize genotypes, consistent with the shared genome. The response to ozone stress differs dramatically in the magnitude of response but many of the responsive genes are shared, suggesting that the observed physiological variability may be attributed to differences in the regulatory component of the genome of these genotypes rather than in their gene content. Further, differential expression is found in genes involved with the photosynthetic machinery of the leaf, and a key element to reduction in sensitivity is a controlled response to the presence of ozone.

## Results

We designed this study to answer the following questions: Is the expressed transcriptome of the leaf part of the shared genome? Are expressed transcripts structurally different between genotypes and conditions (alternative splicing)? Is the transcriptional response to ozone similar among all genotypes, or is there evidence that may explain the different physiological response to ozone in the transcriptome? To answer these questions, we used long read technology and generated more than 6 million consensus reads across ten libraries. This enabled us to move beyond locus level identification and assay actual transcript variants as the fundamental components of the transcriptome definition. A complimentary in-depth study of 120 samples, 12 independent replicates of each condition (5 genotypes, ambient/ozone) using short-read technology enabled precise quantification of the transcriptome.

### Long read transcriptomic data reveal a shared leaf transcriptome

There are a total of ~6 million long reads from Pacific Biosciences Sequel I technology in 10 different samples corresponding to 5 genotypes (B73, Mo17, C123, NC338, Hp301) in ambient and ozone conditions. Do the leaf transcriptomes of different maize lines represent the shared genome, or, on the contrary, reveal the genotype specific genes? Reference genomes are available for the B73 [20] and Mo17 lines (two versions Mo17 Yan [21] and Mo17 Cau [17]). We use IsoSeq3, which does not require a reference genome, (https://github.com/PacificBiosciences/IsoSeq3) to obtain transcript models for each genotype/condition individually. We map all ten transcriptomes to the three available reference genomes. While mapping rates are slightly higher for the percentage of B73 transcripts mapping to B73 and of Mo17 transcripts mapping to Mo17 references, the differences are negligible and C123, NC338 and Hp301 transcriptomes map equally well to all three references (mapping rates 97.5%-99%) (Figure 1A, Supplementary Table 1). There are very small numbers of unmapped transcripts. Of the two Mo17 genomes, the Cau reference have fewer unmapped transcripts than the Yan, suggesting an improvement to the genome assembly (Figure 1A, Supplementary Table 1). The high mapping rates strongly suggest a shared transcriptome.

**Figure 1.**
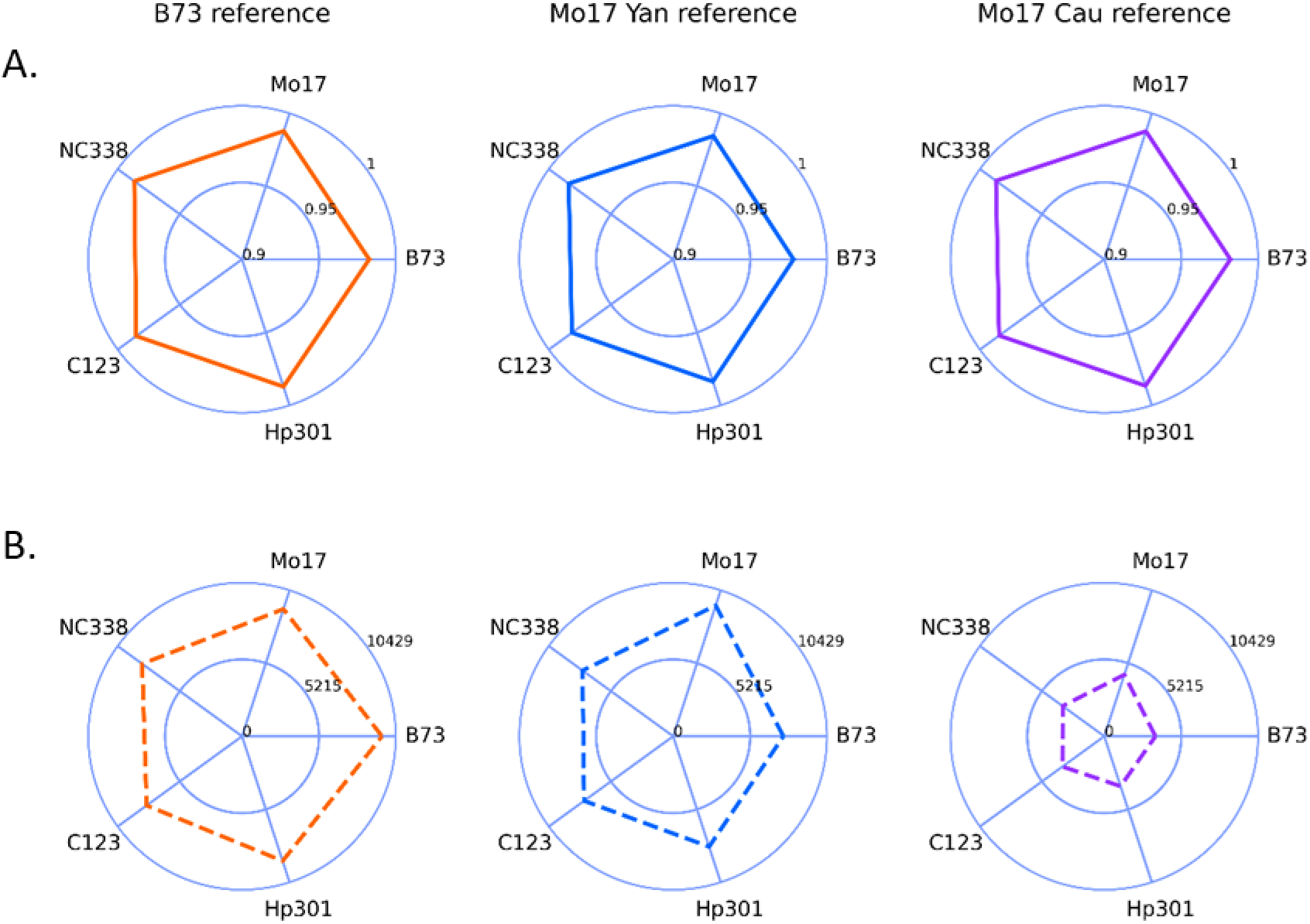
Long read transcriptomes. Each column represents one of the Z. mays reference genomes mapped to a maize reference genome: B73 [20] (left, orange), Mo17 Yan [21] (center, blue), Mo17 Cau [17] (right, purple). Panel A: The proportion of mapped transcripts for each genotype. Each library was processed independently, and the mapped transcripts in both conditions were combined for each genotype. The proportion for each genotype is a point with the estimates connected by a solid line. The internal area can be compared across columns and the similarity in the area is due to the high degree of similarity in the proportion of transcripts for each genotyped mapped for all references. Note the scale ranges from 0.9 to 1.0. Panel B For each genotype/condition the number of full-splice match (FSM) transcripts is identified using SQANTI [22]. The average for each genotype is plotted as a point with the estimates connected by a dashed line. Scale 0 to 10429 transcripts. The internal area is different for each of these with the Mo17 Cau reference the smallest. Data are in Supplementary Table 2.

We use SQANTI to associate transcripts with genes in the reference genomes and quality control [22]. Full length transcripts that match a reference transcript in all junctions are referred to as full-splice matches. Although the absolute number of transcripts obtained for each library varied from 14,000 to 24,000, possibly due to sequencing depth differences, we found a high mapping rate for the full-splice matches regardless of which of the three maize reference genome sequences were used (Figure 1B). There were fewer full-splice matches in the Yan reference, reflecting a lower number of annotated transcripts in this reference, and the highest amount in the B73 refernce due to the greater depth of annotation in this reference (Figure 1B).

To determine whether the transcripts are from orthologous loci in B73 and Mo17 we analyze our data in the context of synteny information using SynMap [23] and SynFind within CoGe [24]. Ninety three percent of the transcripts from B73 and Mo17 map to syntenic loci in the reference genomes. Of the 7% that map to non-syntenic genes, the vast majority of these (80%, 713 out of 903) are consistently expressed in all 5 genotypes. The syntenic list of B73/Mo17 from Sun et al. [17] shows a similar result (94.5% mapping to syntenic loci).

Transcripts that map to genic regions without annotated transcripts (novel in SQANTI) are not part of the syntenic genome lists, as these loci have not previously been annotated. To determine whether these loci are part of the shared genome we looked at both short and long reads. Out of the 1,575 novel, fusion, or antisense loci, 1,572 are expressed in multiple genotypes, indicating these are also part of the shared genome (Supplementary Figure 1A, B). There are 3 novel loci exclusive to B73 with more than 5 short reads in both conditions (Supplementary Table 3). BLAST results of the transcripts within these loci finds partial hits to B73, Mo17, and other cultivars, and are similar to putative transposase and retrotransposon genes, making them unlikely to be unique to B73.

*We do not find evidence supporting the presence of genes expressed in the leaf that diverge in presence/absence variation among the genotypes. All of our data point to a single expressed transcriptome for the leaf that is part of the shared genome, with most loci being syntenic.*

### The majority of genes in the maize leaf transcriptome have a single transcript, with multi-transcript genes having few alternate exons

To estimate the leaf transcriptome for maize we combine transcripts from all genotypes and conditions using chaining (https://github.com/Magdoll/cDNA_Cupcake). The leaf transcriptome contains a total of 48,950 transcripts corresponding to 14,877 genes, with 2,273 novel loci and 12,604 previously annotated genes. 6,895 leaf genes have more than one transcript variant with 17% of the genes expressing more than 3 transcripts. Overall, the average number of transcripts per gene is ~2. 5 and the median is 2.

Splicing alters the number and structure of exons. We quantified exon structure in the B73 reference (Figure 2A) and in the leaf transcriptome (Figure 2B). The number of alternate donors and acceptors (as represented by the number of exon fragments) increases as the number of transcripts per gene increases in the leaf transcriptome (Figure 2B), similar to the B73 reference transcriptome (Figure 2A). However, the number of exon regions remains fairly consistent in the leaf transcriptome, even for genes with greater numbers of transcripts per gene.

**Figure 2.**
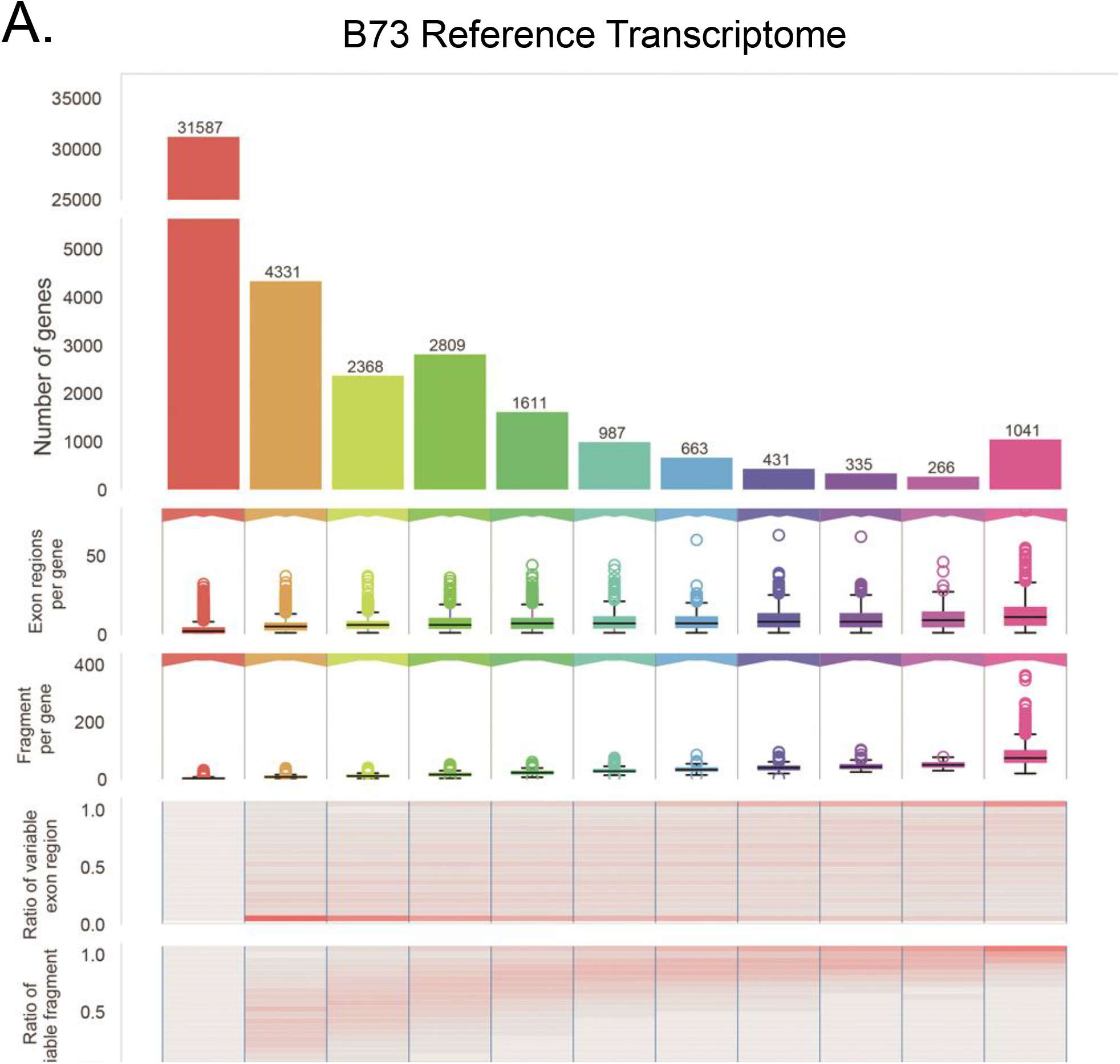
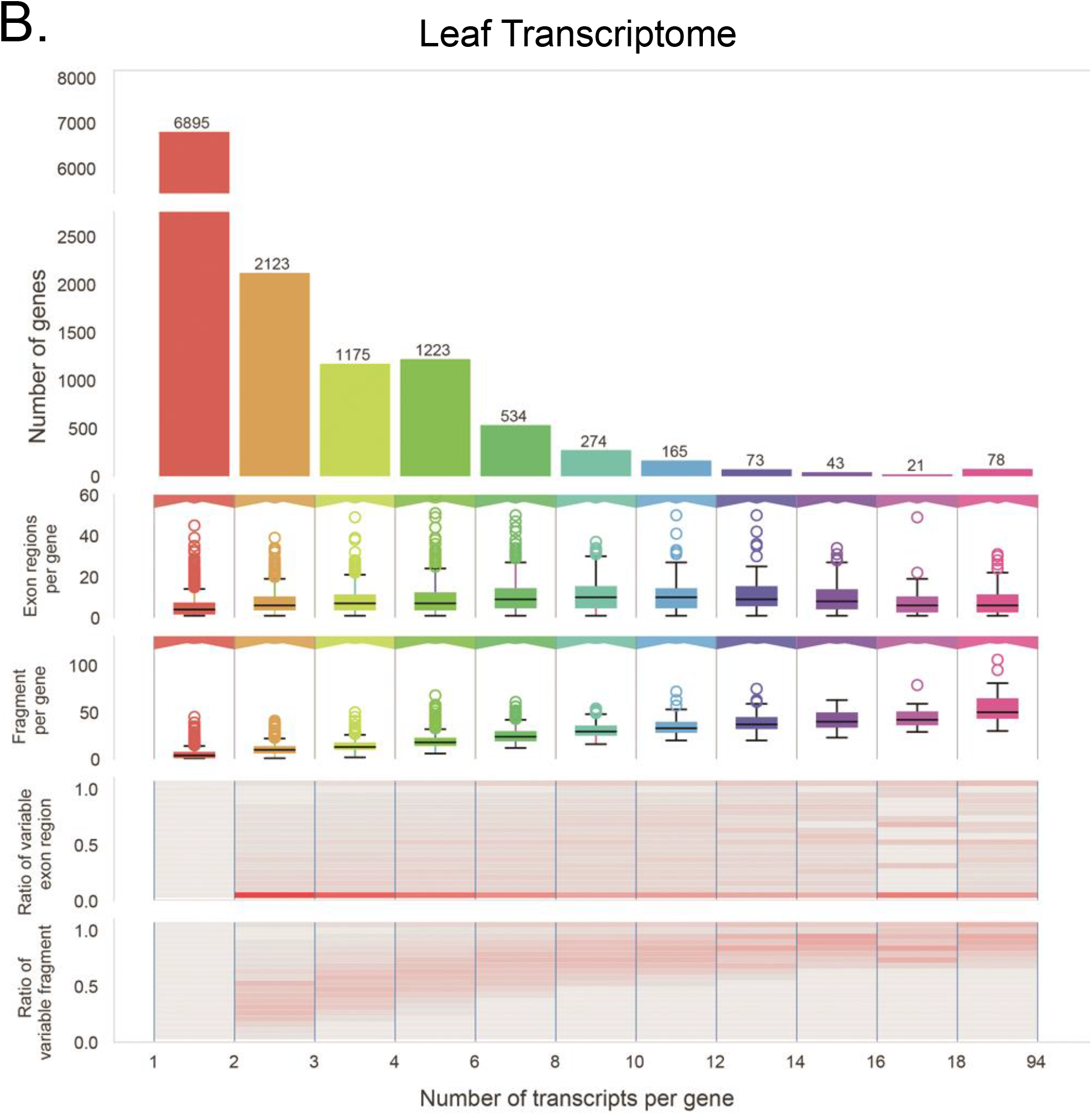
Transcriptome complexity. The x-axis is the number of transcripts. Top row, a bar plot for the number of genes with *x* transcripts. Second row a box and whisker plot for the distribution of the number of exon regions for the genes with x transcripts. Third row a box and whisker plot for the distribution of the number of fragments for the genes with *x* transcripts. Defined as in [66]. Third row, for each transcript the proportion of variable/alternative exons in each gene with *x* transcripts. Fourth row the proportion of variable fragments in each gene with *x* transcripts. that summarize the variation in overlapping exons. Higher proportions of variable exon regions indicate higher proportions of alternative exons and higher proportions of variable exon fragments indicate a higher proportion of alternative donor and acceptor sites. Panel A) all transcripts with unique splice junctions for the B73 reference transcriptome (133,204 transcripts, 46,430 genes) and the (B) Leaf transcriptome from 5 genotypes (B73, C123, Hp301, Mo17, NC338) and 2 treatments (ambient, elevated ozone) (31,388 transcripts, 12,604 genes). B73 leaf transcriptome from [29] (Supplemental Figure 1A).

### Transcripts with intron retentions are lowly expressed

Approximately ~48% of the genes with two expressed transcripts have one transcript with an annotated intron retention (IR). Estimates of the expression of these isoforms with IR are low (Supplementary Figure 1C, D) with, on average, less than 20% of the total gene expression.

### There is limited evidence for alternate splicing in the structure of the leaf transcriptome

Despite reports of splicing as a consequence of response to stress in plants [25–27, reviewed in 28], the leaf transcriptome estimated from long reads in this study have fewer alternate exons than the B73 reference. In order to ensure we have not overlooked potential alternative exons, we compare our estimate of the leaf transcriptome to a previously published long read leaf transcriptome from B73 [29] (Supplementary Figure 2A). These data show a similar estimate of the number of genes expressed (~21%) compared to our estimate (~27%). The distribution of the number of transcripts per gene and the distribution of alternate exons are also similar. Overall, the Wang et al. 2018 B73 leaf transcriptome is slightly less complex with 96% of the genes having 3 or fewer transcripts compared to our finding of ~87%. We expect to have more evidence of complexity with more than 6 million reads from 5 genotypes and 2 conditions, compared to the Wang et al. 2018 B73 study (~550,000 reads in the leaf). This result highlights the critical contribution of deep sequencing for the definition of a transcriptome and suggests that sequencing of multiple replicates is important for transcript annotation.

*Taken together, our results portray a leaf transcriptional landscape that includes more genes than previously described but that is mostly composed of genes with single or few transcript variants, with those variants primarily alternative donor/acceptors rather than alternative exon cassettes.*

### Transcript structure and expression levels are similar among genotypes

We estimate gene expression in ambient and ozone stress conditions for each genotype by mapping Illumina short read data from 120 individual plants (n=12 per experimental condition; n=24 per genotype) to the maize leaf transcriptome estimated from long read data as described above using RSEM [30]. Gene expression is similar in ambient conditions across all genotypes. There is no evidence of presence/absence variation among the 5 genotypes (Supplementary Figure 3). Most genes are consistently expressed in all 10 experimental groups (TPM>5 in 50% of the replicates). Indeed, more than two-thirds of the genes in this study are consistently expressed in all ten genotype/treatment combinations. Consistent with sampling variation [31], genes detected in fewer samples are expressed at much lower levels (Figure 3).

**Figure 3.**
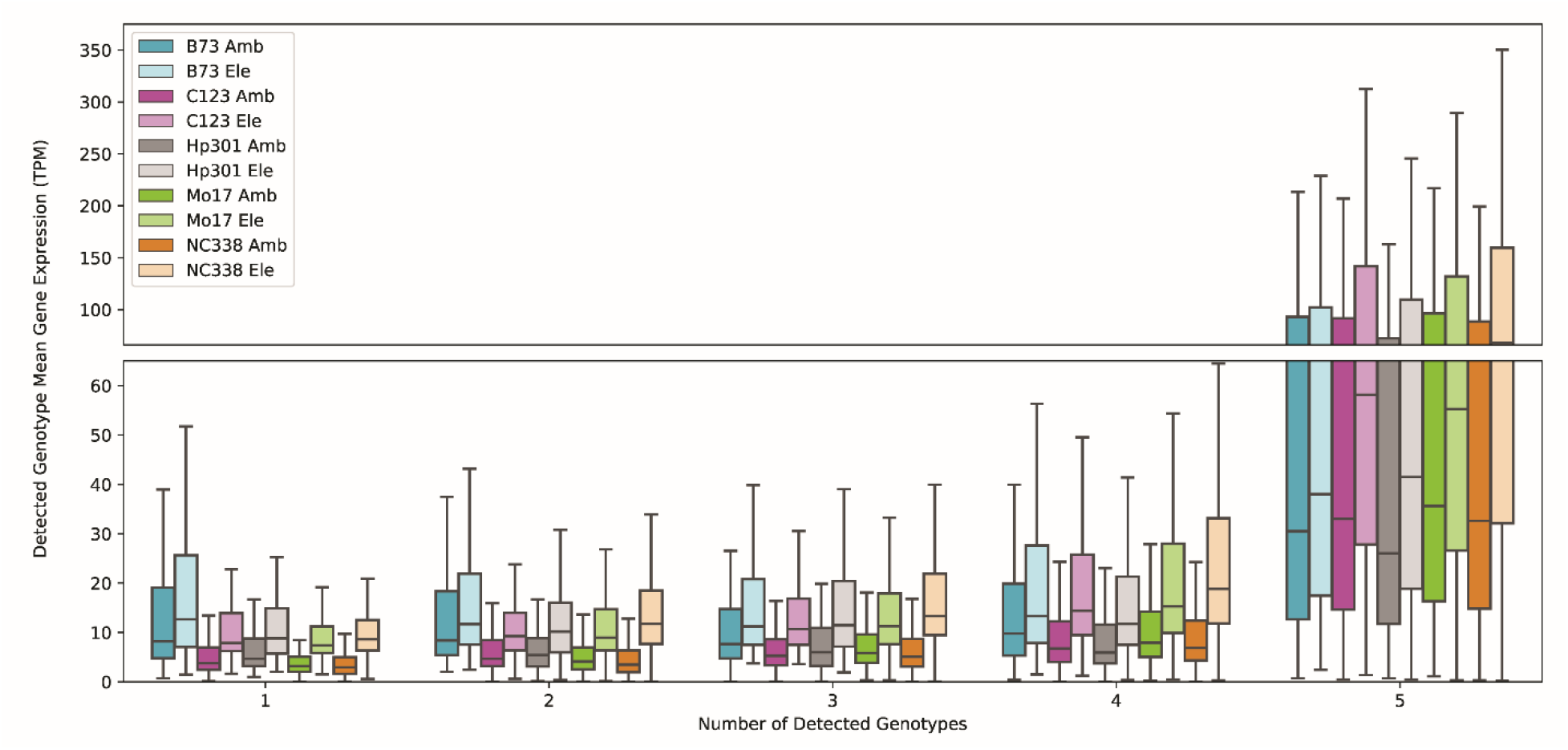
Gene expression. Mean gene expression (TPM) for each genotype and treatment. B73 is blue, C123 is red, Hp301 is gray, Mo17 is green and Nc338 is orange. Transcripts are consistently detected if their TPM values are greater than 5 in 50% of the replicates in either ambient or elevated ozone condition for that genotype. Genes are detected when at least one transcript is detected. For genes detected in 1-4 genotype(s) expression values are low, while for genes detected in all 5 genotypes expression values can be much higher. This indicates that it is unlikely that presence/absence variation is at play, and instead more likely that presence/absence is due to sampling variation [31].

### Alternative splicing is not a major contributor to the maize leaf response to ozone stress

We evaluated differential isoform usage in the transcriptional response to ozone stress. We assayed all genes with multiple transcripts in a particular genotype for differential isoform usage between ambient and elevated ozone conditions (Supplementary Figure 6). 59 genes are detected as having major isoform switching in NC338, and fewer than 35 genes in each of the other 4 genotypes, with 1 gene (Zm00001d043613) having evidence for major isoform switching in B73 (Supplementary Table 5). 18 genes with differential isoform usage have annotation linked to plant stress responses. Many of the genes with differential isoform usage have dramatically different proteins in their annotated isoforms (e. g. Supplementary Figure 7). This suggests that an ozone stress response may switch the proteins produced by these genes, or that there are two similar genes in proximity in the genome annotated as a single gene and that the expression of one of these genes responds to ozone stress, while the other does not respond.

### Genotypes differed in the magnitude of the response to ozone stress

B73 showed a limited response to elevated ozone, with 151 differentially expressed genes, while NC338 showed evidence for more than 3,300 differentially expressed genes (Figure 4A). The number of differentially expressed genes is progressively greater in the other genotypes of Hp301 (1807 genes), C123 (2085 genes), Mo17 (2382 genes) and NC338 (3384 genes). The number of genes differentially expressed in a genotype-specific manner followed the same sequence: B73 (8 genes), Hp301 (157 genes), C123 (223 genes), Mo17 (333 genes) and NC338 (998 genes) (Figure 4A). 789 are differentially expressed in common across the four more ozone sensitive genotypes. 83 genes are identified as differentially expressed in all five genotypes, thus representing the common transcriptional response to elevated ozone concentration. We further compare genes differentially expressed (DE) across genotypes and evaluate the magnitude of their differential expression to better understand similarities and differences among genotypes.

**Figure 4.**
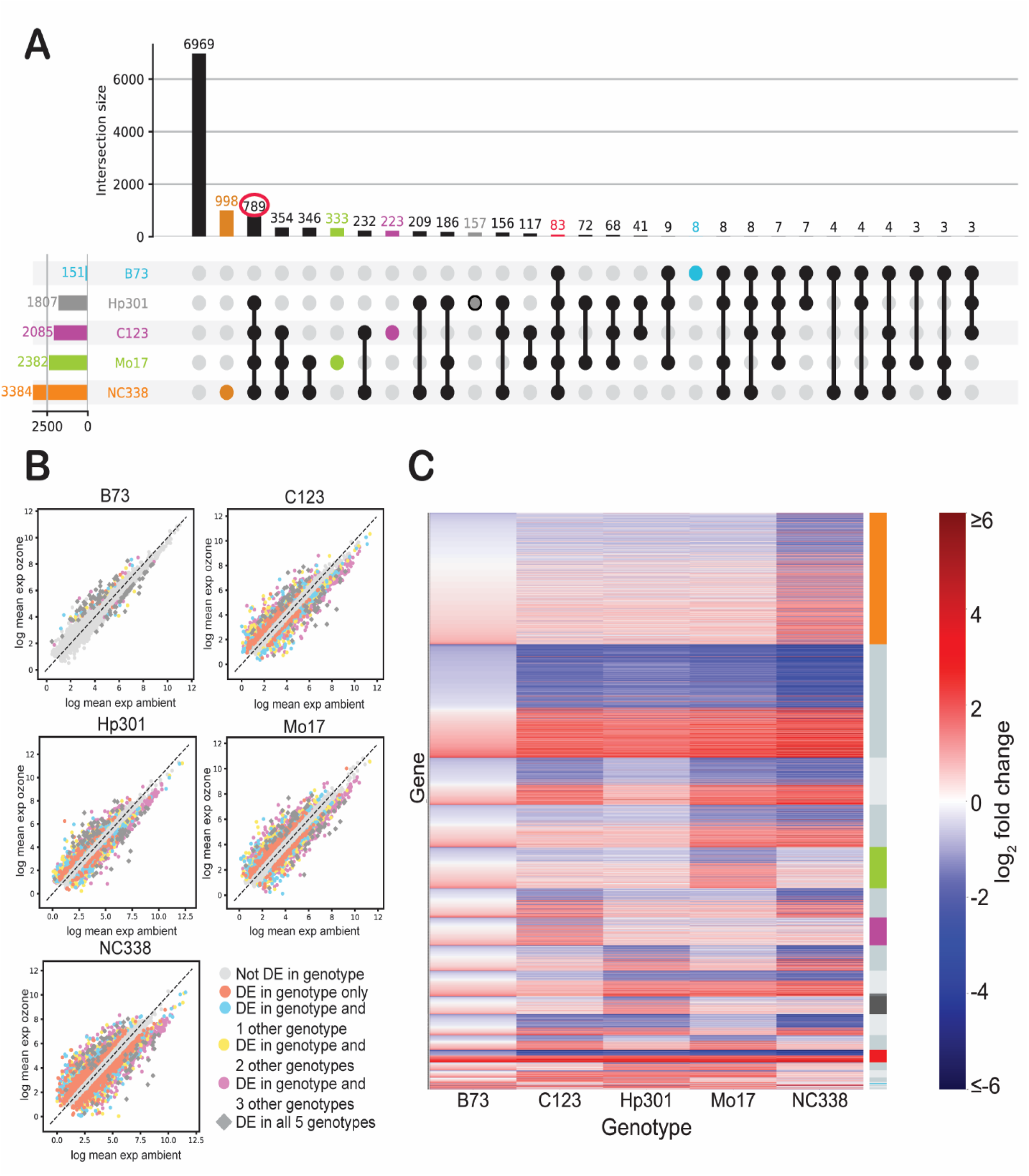
Differentially expressed genes. (A) Many genes are significantly differentially expressed in NC338 only (998, orange) or in all genotypes excluding B73 (789). Relatively few genes are differentially expressed in all genotypes (83, red circle). (B) For each genotype, a scatterplot is shown of the log2(TPM) ambient on the X axis and the log2(TPM) ozone on the Y axis. Differentially expressed (DE) genes in the indicated genotype are in pink, in the indicated genotype and one other genotype are blue, in two other genotypes are yellow, in 3 other genotypes are purple and in all 5 genotypes (gray diamonds). The largest fold changes are consistently apparent win the gray diamonds. Transcripts DE in only one genotype have smaller fold changes than those detected in more than one genotype (as seen by the orange points nearer the diagonal). (C) Genes significant for differential expression. Categories are ordered as in Panel A. Rows are genes and columns are genotypes. B73 has overall lower fold changes than the other genotypes and NC338 has larger fold changes compared to the other more sensitive genotypes. The genes differentially expressed in all genotypes are indicated by the red bar in the right column of the figure and other colors match the colors in panel A.

For genes significant in only one genotype, the estimated fold change is relatively modest, with the B73 response substantially smaller in all comparisons (Figure 4B, 4C). In each genotype, enriched gene sets [32] for the Gene Ontology biological process category included photosynthesis (e. g. photosystem I, protein-chromophore linkages) (Supplementary Table 6, flag_detect_DE_*genotype*).

### Diverse genotypes show a common transcriptional response to ozone stress

Are differential responses to ozone stress associated with gene expression differences?_The largest fold changes are found in genes significant in all genotypes (Figure 4B), and the proportion of up- and down-regulated genes is balanced for all genotypes, including B73 and NC338 (Figure 4B and Supplementary Figure 5). _The regulation of expression in response to ozone is in the same direction for 82 of the 83 transcripts differentially expressed in all 5 genotypes (Figure 4C, red). Zm00001d027525 (basic endochitinase B, EnsemblPlants)) is the only transcript that is down-regulated in B73 under ozone conditions but up-regulated in the other 4 genotypes. Genes differentially expressed in all 5 genotypes are enriched for the following GO categories: biological process (heat shock response, photosynthesis (light harvesting in photosystem I, light reaction, response to red light, and protein-chromophore linkage)); cellular component category (photosystem I, photosystem II and plastoglobule); and molecular function category (chlorophyll binding and pigment binding) (Supplementary Table 5, flag_detect_DE_all5).

In addition to many of the same GO categories above related to photosynthesis, for genes significantly differentially expressed in the four more sensitive genotypes C123, Hp301, Mo17, and NC338 (Figure 4A (n=789)), additional categories are enriched. These include GO biological processes related to metabolic processes (aromatic amino acid family biosynthetic process, carotenoid biosynthetic process, coenzyme biosynthetic process, glycine catabolic process, hydrogen peroxide catabolic process, oxylipin biosynthetic process, regulation of lipid metabolic process, unsaturated fatty acid biosynthetic process), photosynthesis (light harvesting in photosystem I, light reaction, photosynthetic electron transport in photosystem I, photosystem I assembly, reductive pentose-phosphate cycle, triose phosphate transmembrane transport, response to red light, response to light stimulus, response to low light intensity stimulus), protein-chromophore linkage, iron-sulfur cluster assembly, and detection of biotic stimulus (Supplementary Table 6, flag_detect_DE_all_noB73). The direction of regulation of the genes significant in these 4 more ozone sensitive genotypes is in the same direction for 788 of the 789 genes (~46% up-regulated, ~54% down-regulated) with Zm00001d037513 (germ-like protein1, EnsemblPlants) up-regulated in C123 and down-regulated in the other 3 genotypes (Figure 4C).

### Chlorophyll content estimate from a spectrophotometer follows the same pattern as gene expression

The transcriptional response to ozone stress is largely attributable to genes involved in photosynthetic and respiratory functions in the leaf. These general effects are common across genotypes, but the magnitude of response varies. We hypothesize that the magnitude of the transcriptional response to ozone is correlated with changes in leaf chlorophyll content. To evaluate this hypothesis, we re-analyze the ambient and ozone data in Yendrek et al. [33] for the maize genotypes in this study. In agreement with our transcriptional analysis, we found that chlorophyll levels are significantly lower under elevated ozone conditions for NC338 (p<0. 0001), Mo17 (p<0. 001), and C123 (p=0. 003), while there is no significant difference in chlorophyll content in the ambient and elevated ozone for B73.

*In summary, our results indicate that there is a common transcriptional response to ozone stress in the maize leaf. Moreover, the magnitude of the expression response to elevated ozone mirrors physiological differences among maize genotypes.*

## Discussion

Despite the diversity of the maize genotypes and the extensive literature on structural variation among maize genotypes, leaf transcriptomes from Hp301, NC338, and C123 map well to both B73 and Mo17 genomes. A close examination of the few unmapped transcripts points to technical artifacts with no evidence that genes expressed in the leaf have presence/absence variation in any of the genotypes. Indeed, 94.5% of genes expressed in the leaf are part of the annotated syntenic genome. Further, the expression patterns in ambient conditions are similar among all 5 genotypes. This is perhaps surprising given the large degrees of structural variation in the genes [8–11] and estimations of the diversity of maize genomes [19], but is consistent with the decades of mapping work that leveraged markers to map recombinant inbred lines across wide crosses [34], and with the recent Mo17 genome assembly [17] which reported presence/absence variation in only 122 genes between B73 and Mo17. Gene expression in the leaf is limited to the shared genome consistent with a single domestication event giving rise to all maize landraces [35] and with the similarity of leaf tissue between maize and sorghum [29].

The pattern of the expression response is very similar in loci differentially expressed in all 5 genotypes (82/83) and in loci differentially expressed in NC338, Hp301, Mo17 and C123 (786/787), even as the magnitude of expression in response to elevated ozone is dramatically different. We see no evidence for structural variation in the genes that drive the expression response to ozone but cannot rule out variation in *cis* regulation of these genes, which may well be influenced by adjacent variable genomic content [11, 36–39].

There are 83 differentially expressed genes in response to elevated ozone in all 5 maize genotypes. Decreased expression of genes involved in light harvesting dominated the conserved transcriptional response. These include chlorophyll biosynthesis genes, chlorophyll a/b binding proteins, and subunits of photosynthetic proteins. Down-regulation of photosynthesis is a common transcriptional response to ozone [40], and chlorophyll biosynthetic genes are also reduced in an ozone sensitive Medicago accession [41]. In an ozone sensitive soybean genotype, decreased photosystem II gene expression is observed after 24 hrs of ozone exposure [42]. The sensitivity of photosynthesis to ozone is clear across both C_3_ and C_4_ crops, and remains a critical target for improving ozone tolerance [43]. Perhaps to combat damage to photosynthetic proteins, increased expression of heat shock proteins is another common response of the five genotypes to elevated ozone.

The genotypes assayed here have been shown to differ in photosynthetic capacity under elevated ozone [6] and they differ in transcriptional responses to ozone stress. Allelic variation in transcriptional response to stress has been documented [44]. Over 20 times more genes are differentially expressed in NC338 in ambient and elevated ozone compared B73, and the transcriptional response of B73 is muted relative to the other genotypes. In a companion field experiment, there is a greater decline in chlorophyll content associated with accelerated senescence in inbreds C123 and NC338 as compared to B73 [33]. The metabolite profile of B73 was not responsive to elevated ozone in the field [45]. Additionally, hybrids containing parents Hp301 and NC338 showed greater reductions in photosynthesis under elevated ozone in the field [7]. Ozone tolerance has been associated with a dampened transcriptional response in genotypes of other species including Arabidopsis [40], soybean [42, 46] and Medicago [41]. A dampened transcriptional response to other abiotic stresses including drought [47], cold stress [48] and salt stress [49] was observed in tolerant genotypes.

These results are consistent with oxidative stress tolerance being associated with a weaker response to a reactive oxygen species signal. Notably, B73 has equivalent, or greater, stomatal conductance than the more sensitive maize genotypes studied here [33]. This means that the flux of ozone into the leaf mesophyll would tend to be greatest in B73. That an equal or greater ozone dose elicited a weaker transcriptional response, suggests that genetic variation in downstream processes is key to ozone tolerance i.e. the processes that either sense ROS in the apoplast, transduce signals, produce the secondary burst of ROS in the cytoplasm, or respond to the secondary ROS burst e.g. plant growth regulator mediated pathways [50]. For example, GO enrichment analysis (Table S5), indicated differential expression of genes related to *cellular response to hydrogen peroxide*, *detection of biotic stimulus*, *hydrogen peroxide catabolic process*, and *regulation of response to biotic stimulus* in the ozone sensitive genotypes, but not B73. At the same time, there is differential gene expression related to *response to salicylic acid* in B73 that is not observed in the more ozone sensitive genotypes. The strength of ozone impacts on Arabidopsis varies with the strength of ethylene-signaling components of the pathway when modulated genetically or chemically [51]. The transcriptional data presented here suggest a similar strategy might be relevant in maize and could assist in the identification of potential targets for manipulation.

## Conclusions

Maize genes underlying the expressed transcriptome of the leaf are part of the syntenic genome. There is a conserved response to ozone exposure among 5 maize genotypes, but the extent of the response varied. We find that the leaf transcriptomes for all 5 genotypes are from the syntenic genome (94.5%) and are conserved in transcriptome content and expression in ambient conditions. Transcripts not expressed consistently in all genotypes and conditions showed low expression, suggesting that sampling variation rather than presence/absence variation is responsible for this observation. While the overall response to ozone is largely consistent, the extent of the response is dramatically different among the genotypes, ranging from 150 genes significantly differentially expressed (B73) to more than 3,000 genes significantly differentially expressed (NC338). B73 has a dampened transcriptional response, suggesting potential fundamental differences compared to the other 4 genotypes.

We found no support for structural variation in the expressed genome of the leaf but did find that all genotypes show regulatory responses in genes related to photosynthesis, while the extent of that response varies. Our findings show that ozone directly impacts the photosynthetic machinery of the leaf, and a key element to reduction in sensitivity is a controlled response to the presence of ozone.

## Methods

### Experimental design and sample collection

Twenty-four plants from each genotype were grown in 5. 7 L pots in LC1 mix (Sun Gro Horticulture Distribution Inc., Bellevue, WA, USA) in six growth chambers (Growth Chamber, Chagrin Falls, OH, USA) set to maintain a 15 hr day at 25 °C and photosynthetic photon flux (PPF) of ~350 □mol m^-2^ s^-1^, 9 hr night at 21 °C, and a relative humidity of 60%. We grew 120 plants in the six growth chambers, with three chambers maintained at low ozone conditions (< 10 ppb) and three at elevated ozone conditions (100 ppb). Ozone is generated using a variable output UV-C light bulb ballast (HVAC 560 ozone generator, Crystal Air, Langley, Canada) and plants are fumigated with 100 ppb ozone for 6 hrs per day from emergence to sampling. Four plants of each of the five genotypes, B73, C124, Hp301, NC338, Mo17 were randomly assigned to each chamber for a total of twenty-four plants per genotype. Plants were fertilized (Osmocote Blend 19-5-8) at the start of the experiment. Twenty-six days after planting, leaf tissue from the 5^th^ leaf is sampled directly into liquid N_2_ for a total of 120 independent leaf samples (5 genotypes x 4 plants per genotype x 2 treatment conditions x 3 growth chambers per treatment) for a total of 12 replicates per genotype and ozone treatment. Leaf samples were sent to University of Florida where leaf punches were collected in a cold room on dry ice. A separate sample from a single plant from each genotype and ozone treatment is collected for long read sequencing.

### Long-read library preparation and sequencing

Leaf tissue (~175 mg) from a single individual plant of each genotype/treatment combination (n=10) was submitted to the University of Florida Interdisciplinary Center for Biotechnology Research for RNA extraction, library synthesis and long-read sequencing. Libraries were sequenced and processed on the PacBio SEQUEL platform using 1 LR SMRT cell per sample and SMRT Link 6.0.0 chemistry. The Mo17 ambient library was sequenced twice, once to optimize sequencing parameters and again when the other 9 libraries were sequenced. Data have been deposited at the SRA (BioProject accession PRJNA604929).

### Long-Read processing

For the primary data collected as a part of this study, the IsoSeq3 pipeline (v3.0.0) is applied to each library separately. CCS (v3.1.0) is used to estimate consensus reads. Primers are removed using lima (v1.7.1). IsoSeq3 cluster is used to trim polyA tails, remove false concatemers and cluster full-length reads by sequence similarity. Consensus (‘polished’) sequences for each cluster are generated using IsoSeq3 polish. The expected error rate after this step is less than < 2% [52]. Clustering and polishing of trimmed reads yielded 24,000 to 44,000 high quality individual transcripts per library.

### Mapping long reads to multiple maize genomes

For the primary data collected in this study polished transcript long reads are mapped to reference genomes B73 (Ensembl version 41, [20]), Mo17 Yan [21] and Mo17 Cau [17] using minimap2 (v2.12, [53]).

### Estimating the Transcriptome

SQANTI identifies representative isoform sequences. The PacBio data may contain multiple sequences mapping to the same reference transcript. We used_SQANTI (sqanti_qc.py, [22]) to evaluate the transcripts generated from individual libraries and all libraries combined mapped to references (B73 or Mo17 Yan). RNA-seq short reads are mapped to transcripts with the program STAR (2.7.0b, [54]) and junction coverage estimated. Expression is estimated using RSEM (version 1.2.28, [30, 55]). Transcripts with insufficient short-read support, non-canonical junctions, or evidence of a possible reverse transcriptase switching event are noted and removed from further analysis [22].

### Annotation of maize transcripts

FASTA files of translated transcript sequences are used as queries to obtain putative functional, structural or signaling motif information from InterProScan (version 5.29-68.0, [56]), TMHMM (version 2.0, [57, 58]) and SignalP (version 4.1, [59, 60]). Translated transcripts are also linked to proteins in public databases via sequence similarity (see Supplementary Methods for a list of public databases used). Repetitive regions are annotated using RepeatMasker (version 4.0.5, http://www.repeatmasker.org) and putative RNA regulatory sequences are identified using ScanForMotif [61]. Default parameters are used. We collated all the above annotation into a GTF-like file compatible with tappAS [62] (Supplementary File 1).

### Synteny

SynMap [23] and SynFind within CoGe [24] are used to identify collinear gene-sets between the B73 reference and the Mo17 Cau reference (Margaret Woodhouse, maizeGDB).

### Secondary Data

The raw data (.bas.h5) [29] are obtained from the authors. These data are B73 with the following tissue types: leaf, silk, pericarp, bract, shoot and seedling. Data are processed as described above using the B73 reference genome.

### RNAseq library preparation and sequencing

Approximately 20mg of flash frozen leaf tissue from each sample is placed into individual mini tubes in a 96-tube plate format (Axygen MTS-11-C-R). Sample freeze-drying, mRNA purification, cDNA synthesis and library preparation are carried out by Rapid Genomics (Gainesville, FL) to generate an individual dual indexed library per sample. Individual libraries are quantified and pooled to generate equimolar samples for sequencing. The final pooled library is sequenced on 3 lanes of an Illumina HiSeq 3000 at Rapid Genomics and 1 lane of Illumina Novaseq at Novogene (Sacramento, CA). Data have been deposited at the SRA (BioProject accession PRJNA604929).

### Quantification and differential expression

FASTQ files are trimmed of adapter sequence (cutadapt, version 2.1, [63]) and merged if R1 and R2 reads overlapped (bbmerge.py from BBMap version 38.44, [64]). Duplicate reads are removed, and gene and transcript expression values are estimated using RSEM [30, 55] using the estimated transcriptome GTF file (Supplementary File 2). Transcripts are considered consistently detected if their TPM values are greater than 5 in 50% of the replicates in either ambient or elevated ozone condition (Supplementary File 3). For each genotype, transcripts consistently detected in either ambient or elevated ozone conditions are evaluated for differential expression and differential splicing using tappAS (version 1.0.6, [62]) at FDR level [65] of 0.05 as a threshold to declare statistical significance. GO enrichment is performed using Fisher’s exact test and an FDR correction of 0.05 is used as a threshold.

## Acknowledgments

NSF Plant Genome Research Program (PGR-1238030; ADBL, LMM, and EAA) MCA-PGR: Genetic and genomic approaches to understand and improve maize responses to ozone NIH R01GM128193 (LMM), R03CA222444 (AC, LMM) USDA SoyFACE Global Change Research. Project Number: 5012-21000-030-17-S. Brad Barbazuk for helpful discussion, Margaret Woodhouse (with maizeGDB for synteny file), Junping Shi for providing the synteny list from Sun et al. 2018 Deborah Morse and James Resnick for assistance with sample processing. Francisco Pardo-Palacios and Pedro Salguero Garcia for help with maize functional annotation. Dr. Bo Wang, Peter van Buren and Dr. Doreen Ware for graciously sharing maize PacBio raw data.

## Data Availability

Raw data (BAM files for the PacBio data and FASTQ files for the RNA-seq data) are deposited under SRA BioProject accession PRJNA604929. All processed data are attached to this manuscript, including the GTF file for the transcriptome (Supplementary File 2), quantified expression data for analysis (Supplementary File 3) and annotations for the transcriptome (Supplementary File 1). All scripts and full detailed documentation for the analysis are posted on github (https://github.com/McIntyre-Lab/papers/tree/master/nanni_maize_2021).

## Author contributions

**Adalena V. Nanni** developed all analysis plans, analyzed all short-read data, contributed to analysis of long read data, annotated the maize transcriptomes, managed all references, designed and contributed to figures, contributed to the writing of the paper. Worked on the github page.

**Alison M. Morse** designed and executed the analysis performed all long-read analyses, interpreted the findings, designed and contributed to figures, contributed to the writing of the paper and is responsible for the github page.

**Jeremy R. B. Newman** participated in analysis, designed and contributed to figures, contributed to the writing of the paper.

**Nicole E. Choquette** identified genotypes for testing, performed the growth chamber experiment and collected tissue.

**Jessica M. Wedow** identified genotypes for testing, performed the growth chamber experiment and collected tissue.

**Zihao Liu** created the transcriptome visualization plots.

**Andrew D. B. Leakey** contributed to the overall design of the umbrella project (PGR-1238030), interpreted the findings, designed figures, interpreted the findings, contributed to the writing of the paper.

**Ana Conesa** contributed to the analyses of long reads, interpreted the findings, contributed to the writing of the paper.

**Elizabeth A. Ainsworth** contributed to the overall design of the umbrella project (PGR-1238030), designed this particular experiment, identified genotypes for testing, supervised tissue collection, participated in analysis, designed figures, interpreted the findings, contributed to the writing of the paper.

**Lauren M. McIntyre** contributed to the overall design of the umbrella project (PGR-1238030), designed this particular experiment, designed and supervised the analysis, interpreted the findings, contributed to the writing of the paper.

none of the authors have any competing interests

